# Epstein-Barr virus and *Helicobacter pylori* Coinfection Drives Gut and Brain Barrier Dysfunction

**DOI:** 10.1101/2025.05.20.655045

**Authors:** Vaishali Saini, Siddharth Singh, Buddhadeb Baral, Pramesh Sinha, Anamika Singh, Raphael Gaudin, Hamendra Singh Parmar, Hem Chandra Jha

## Abstract

Biological barriers, including blood-brain barrier (BBB) and gut epithelial barrier, are essential to maintain homeostasis and provide protection against pathogenic stress. Here, we investigated the impact of *Helicobacter pylori*-Epstein Barr Virus coinfection on gut-brain axis integrity. Our results demonstrate that coinfection induces significant disruptions in tight and adherens junction (TJ and AJ respectively) proteins in gastric epithelial cells, including downregulation of ZO-1 and E-cadherin. The secretome from these coinfected cells significantly compromises the integrity of BBB-derived endothelial cells, including cleavage of VE-cadherin, loss of TJ proteins such as Claudin-5, and enhanced permeability. Furthermore, the sole exposure of mice to coinfected secretome lead to the loss of cerebral VE-cadherin and ZO-1 and the induction of neuroinflammation, characterized by elevated pro-inflammatory cytokines, TNF-α, amyloidogenic accumulation, and neurocognitive deficits. The secretome driving such forces potentially carries pathogenic metabolites, host inflammatory cytokines and metabolites that might contribute to disease pathogenesis as observed in this study. Together, our current study provides novel insights into how coinfections contribute to gut-brain axis dysregulation and neurological pathogenesis.

Graphical Abstract

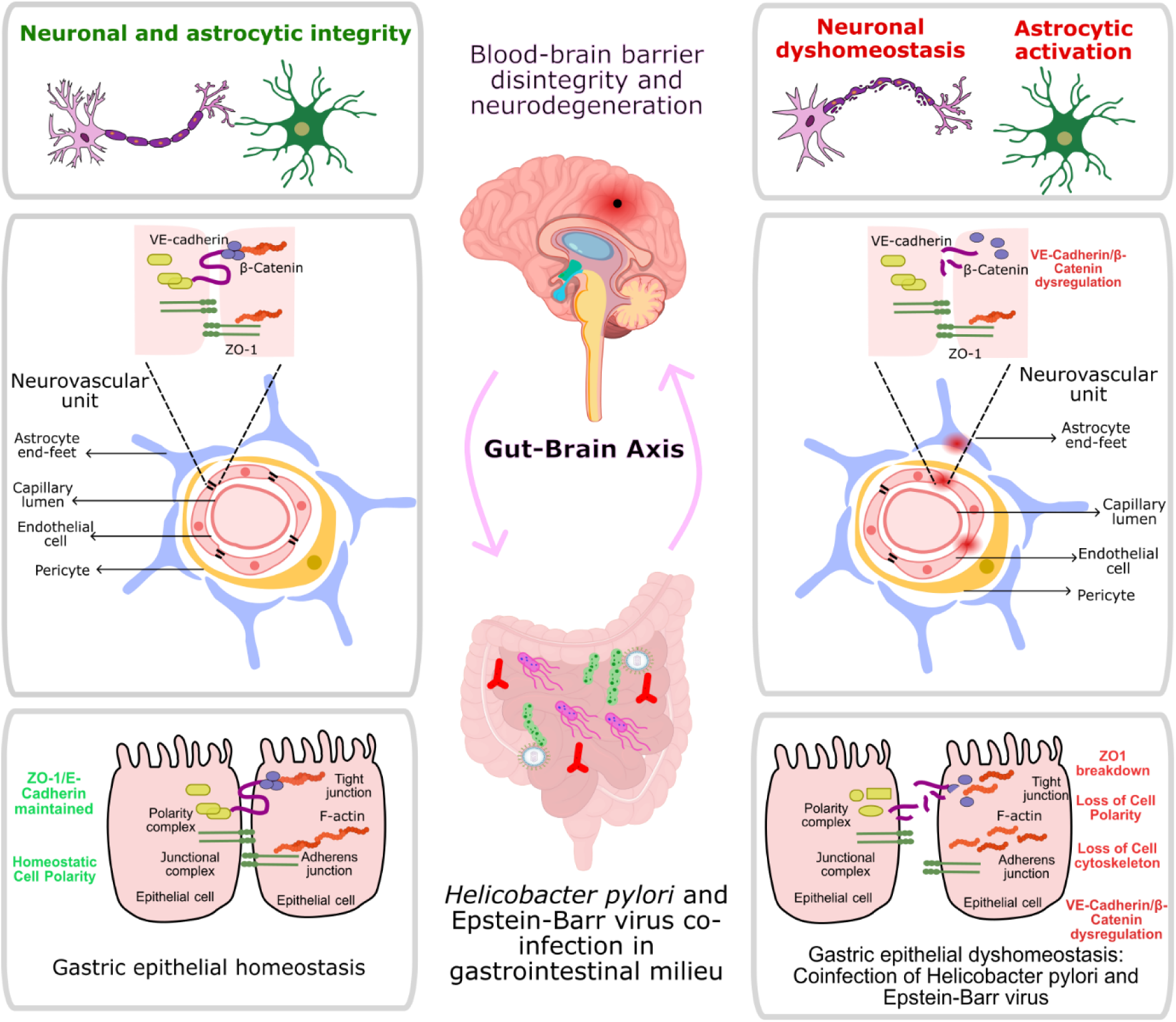

## Introduction

Biological barriers, including the blood-brain barrier (BBB) and the gut barrier, serve as critical interfaces for the transduction of environmental signals. Both barriers regulate influx and response to external stimulus, thereby maintaining homeostasis and protection against pathogenic stress. Integrity of both the barriers is dependent upon structural alterations majorly governed by various environmental cues. Such environmental cues that can cause negative alterations include pathogens such as viruses, bacteria, parasites, toxins, pollutants, and poor nutrition(*1*). Intestinal microbiotas thrive in a mutually beneficial relationship with the human body(*2*, *3*). Microbial dysbiosis can cause barrier disruptions, particularly in BBB. Negative influences on gut and brain barriers can exert long-lasting cognitive consequences as loss of barrier integrity can be plausibly implicated in disease pathogenesis. The intestinal epithelial barrier comprises epithelial cells sealed together with tight junctions. This layer permits the travel of substances *via* transcellular and paracellular pathways. Gastric lumen harbors a plethora of microbiota, including pathogenic microorganisms(*4*). Disruption of this epithelial barrier can plausibly lead to systemic inflammation and translocation of gut microbiota or metabolites to extra-gastric regions such as the central nervous system (CNS). The CNS is protected by specialized endothelial cells that are closely synchronized with pericytes, astrocytes, and neurons, forming the neurovascular unit (NVU). The macromolecular complexes and specialized transport systems of the NVU are responsible for the extremely low transcellular and paracellular permeability of the BBB(*5*). Therefore, impairment of the functions of the brain endothelial cells is considered one of the earliest events leading to severe neurological deficits. Multiple pathogenic metabolites are known to modulate the NVU, leading to neuropathological consequences(*6*).

Gut microbiota plays a critical role in maintaining gut homeostasis. Although majorly composed of bacteria, gut virome and its association with different microorganisms can be fundamental for regulating gut homeostasis. *Helicobacter pylori* (*H. pylori*) and Epstein-Barr virus (EBV) are two major pathogenic agents in the gut, with a propensity of 80% and 10% to induce gastritis and leaky gut symptoms(*7*). This viral-bacterial interaction and pathogenesis can contribute to the progressive loss of gut barrier function. A crosstalk between these pathogens could induce chronic inflammation, the coinfections have been implicated in non-malignant disorders. *H. pylori* is a gram-negative bacterium that colonizes the stomach microenvironment and induces persistent infection through the expression of certain proteins such as cytotoxin-associated gene A (CagA), blood-group antigen-binding adhesion (BabA), sialic-acid binding adhesion (SabA), etc(*8*, *9*). On the other hand, EBV is a gammaherpesvirus that primarily infects epithelial cells and circulates further through B-lymphocytes, ultimately establishing a lifelong latent infection(*10*). Coinfection of both these pathogens induces deleterious inflammation and tissue damage with rise in pro-inflammatory cytokines. For instance, during infection, *H. pylori* contributes to cytokine storms such as interferon γ (IFN-γ), interleukin-6 (IL-6), and IL-13, which further promote EBV proliferation and even higher levels of cytokines such as IL-1β, tumor necrosis factor α (TNF-α) and IL-8(*11*, *12*).

A highly regulated epithelium is maintained by apicobasal polarity, organized cytoskeleton, and junctional complexes exhibiting anti-metastatic properties. Lateral cell-to-cell contacts include tight junctions (TJs) such as ZO-1 and adherens junctions (AJs) such as E-cadherin. TJs such as ZO-1, ZO-2, and ZO-3 interact with scaffolding proteins such as occludins and claudins and other cytoplasmic proteins such as Cdc42, PKCζ, etc(*13*). Whereas cadherins such as E-cadherin interact with β-catenin to maintain the actin cytoskeleton integrity of epithelia. Pathogens can disrupt the gut barrier by depolarization of epithelial cells(*14*). For instance, *H. pylori* infection leads to the disruption of TJ proteins of the epithelial barrier such as ZO-1 and JAM, PAR1/MAPK by VacA, CagA, or Urease-dependent activities, inducing EMT-like phenotype. Interference of AJ proteins such as E-cadherin by CagA causes alterations in PAR1 kinase, MMPs, etc(*15*). Additionally, EBV leads to epithelial barrier disruption through repression of E-cadherin in its latency phase(*16*). The viral infection also causes activation of cdc42 and Par6, furthering the loss of epithelial polarity(*17*). Such regulations exist in the BBB as well. Recent studies indicate that BBB leakage, as a result of cortical microbleeds in humans, contributes to Alzheimer’s disease (AD). Adhesion of brain endothelial cells is maintained by an integral membrane protein vascular endothelial cadherin (VE-cadherin) consisting of five domains, one short transmembrane domain, and a cytoplasmic domain further interacting with catenins such as β-catenin. Cleavage of VE-cadherin might contribute to the reorganization of AJs, thereby inducing endothelial dysfunction(*18*). Additionally, TJ proteins such as ZO-1 and claudin-5 are key structural components in the maintenance of BBB integrity(*19*). Due to the central role of the BBB in initiating pathogen-induced neurological complications, it becomes crucial to explore the gut-brain axis modulations in coinfections of two major gastric pathogens, i.e., *H. pylori* and EBV.

Here, we identify the role of TJ and AJ proteins in initiating leaks into the epithelial and endothelial barriers of the gut and brain, respectively. Upon coinfection of *H. pylori* and EBV to the gastric milieu, we have identified potential contributors to gut-brain axis disruptions through cellular barriers. Our data provides evidence for the alteration of TJ and AJ in gastric cells upon coinfection. Exposure of brain endothelial cells to co-infected gastric secretome shows decreased BBB integrity and significant neuroinflammation. Additionally, mice exposed to co-infected gastric secretome exhibit enhanced junctional BBB disruption and neurological loss as evidenced by protein markers, amyloidogenic accumulation, and neurobehavioral loss.

## Methodology

### 1. Cell culture

The human gastric epithelial cell line (AGS), human neuroblastoma cell line (IMR-32), and human glioblastoma cell line (U-87MG) were procured from the National Centre for Cell Science (NCCS, Pune, India). Immortalized human cerebral microvascular cell line (HBEC-5i) was procured from the American Type Culture Collection (ATCC, Rockville, MD). Gastric, neuronal, and glial cells were cultured in Dulbecco’s Modified Eagle’s Medium (DMEM, Gibco, USA) supplemented with 10% fetal bovine serum (FBS, Gibco, USA), 50 U/mL-100 μg/mL Penicillin-Streptomycin solution (Gibco, USA). HBEC-5i cells were cultured in DMEM-F12 media (Himedia, Mumbai, India) with 40 μg/mL endothelial growth supplement (ECGS, Sigma, Massachusetts, USA) and previously mentioned conditions(*20*).

### 2. EBV culture and isolation

HEK293T cell line with stably transfected bacterial artificial chromosome (BAC)-green fluorescent protein (GFP)-EBV (BAC293T) was provided as a kind gift from Dr Erle Robertson (University of Pennsylvania, USA). The cells were grown in the previously mentioned composition cDMEM with 5% FBS and 0.1 μg/mL puromycin (Gibco, USA). Further, cells were induced for 4-5 days with 3mM butyric acid and 20 ng/mL tetradecanoyl phorbol acetate (TPA) (Sigma, USA), and the concentrated virus was obtained through ultracentrifugation (Beckman-Coulter) at 23,500 rpm and 4°C for 90 min(*21*).

### 3. Bacterial culture

The *H. pylori* clinical strain (HJ9) was isolated from the gastric juice of suspected gastritis patients as per the previously mentioned protocol(*1*). Briefly, bacteria were cultured in Brain Heart Infusion broth (BD-DIFCO, USA) supplemented with 10% FBS and selective antibiotics, namely vancomycin, cefsulodin, amphotericin, and trimethoprim at concentrations 10, 5, 5, and 5 mg/mL respectively (Sigma Aldrich, St Louis, MO, USA). The bacterial cultures were maintained in a microaerophilic environment (85% N_2_, 10% CO_2_, and 5% O_2_) at 37°C within an anaerobic chamber (Whitley DG250).

### 4. Coinfection of EBV and *H. pylori* to AGS cells and conditioned media collection

The AGS cells were seeded in a 6-well plate at a density of 0.25 × 10^6^ cells. Upon reaching a 40-50% confluency, cells were infected at an MOI of 100 and 2.5 for *H. pylori* and EBV, respectively. As previously reported, pre-infection of *H. pylori* and a subsequent EBV infection to AGS cells indicated higher pathogenesis and susceptibility towards disease progression; therefore, we adapted a similar infection model in our study(*7*). After 30 h of infection was completed and conditioned media was collected. The cell debris was removed through a centrifuge at 1000 rpm for 5 min with subsequent filtering with a 0.2-micron syringe filter. The conditioned media (CM) from different samples will be further referred to as Uninfected AGS cells conditioned medium (AGS-CM), *H. pylori*-infected AGS conditioned medium (HP-CM), EBV-infected AGS conditioned medium (EBV-CM), and coinfected AGS conditioned medium (CI-CM), respectively.

### 5. Dynamic *in vitro* BBB model

To mimic blood-brain barrier structure in vitro, we utilized a co-culture method in transwell (0.45 μm, Corning, USA) wherein the upper layer (coated with 0.1% gelatin) was seeded with HBEC-5i endothelial cells, meanwhile bottom layer included a co-culture of astrocytic U-87 MG and neuronal IMR-32 cells. Briefly, cells where endothelial cells were seeded separately to the neural co-cultured cells. After 1 day of seeding cells, the media was changed, and transwell was added to the 6-well plate. This induced an *in vitro* BBB model for studying barrier alterations in our current study.

### 6. In vivo

Adult Swiss Albino male mice weighing 25-35 g with an approximate age of 2 months were housed in polypropylene cages and were acclimatized in a 12 h/12 h light-dark cycle before experimentation. The temperature was maintained at 23 ± 2 °C with controlled humidity and *ad libitum* access to food and water. The study was carried out strictly following the regulations on animal experiments as per the Institutional Animal Ethics Committee (IAEC), the Committee for the Purpose of Control and Supervision of Experiments on Animals (CPCSEA), and the Ministry of Environment and Forests, the Government of India. Protocols were approved by IAEC of the School of Biotechnology, Devi Ahilya Vishwavidyalaya, Indore (registration no. 779).

Five groups (n=6) were formed: immunosuppressed control, AGS-CM exposed, HP-CM exposed, EBV-CM exposed, CI-CM exposed, and neat control (negative control). Mice were immunosuppressed with an intraperitoneal injection of cyclosporin (working concentration = 10 mg/kg)(*22*). Mice were subsequently maintained for 10 CM doses, and behavioural assays were recorded up to 4 weeks before sacrifice.

### 7. Neurobehavioral assessments

#### a. 8-arm radial maze (RAM) test

To determine the status of accelerated brain senescence induced by CM treatment, cognitive function was tested in mice by assessing performance in a radial arms maze test. The maze consisted of 8 arms, with each arm 50 cm long × 10 cm wide × 40 cm tall, placed in a dimly lit room with temperature maintained at 23 ± 2 °C. Mice were familiarized or habituated for 3 days before the training period. They were placed in the maze for approximately 10 min with rewards in specific arms during the training period chosen in a pseudo-random fashion. During the experimental phase, the unblocked arms were baited with food pellets, and the mice were allowed to explore for 5 min. After each test, the maze was wiped with 70% ethanol and allowed to dry completely. Two types of errors were recorded: working error, indicating the re-entry of mice in an arm that has been baited and visited, and reference error, indicating entry into an arm that has been previously baited during the training phase.

#### b. Elevated Plus maze (EPM) test

Anxiety-like behavior in mice was tested using the EPM test. The apparatus consists of two open (50 x 12 cm) and two closed arms (50 x 12 x 50 cm) with a central square (12 x 12 cm) in between and elevated 50 cm above the ground. Each mouse was located at the central platform, and its activity was recorded for 2 min. The number of times and the duration for which mice entered the closed and open arms indicated anxiety and less/no anxiety-like behavior, respectively.

### 8. Quantitative Real-Time Polymerase Chain Reaction (qRT-PCR)

Total RNA was isolated using the TRIzol method, as mentioned previously(*23*). Briefly, cell pellets were washed with 1x PBS and RNA was extracted using TRI reagent (Sigma–Aldrich, St. Louis, USA) and was reverse transcribed using PrimeScriptTM RT kit (Takara, Shiga, Japan) according to the manufacturer’s protocol. qRT-PCR was performed using SYBR green real-time master mix (Agilent Technologies, Santa Clara, USA) programmed at 10 min at 95 °C followed by (15 s at 95 °C, 20 s at 58 °C, 20 s at 72 °C) up to 40 cycles. The primer sequences are listed in **Suppl Table 1**. Melt curve analysis was performed to confirm the specificity of PCR amplicons. Human glyceraldehyde 3-phosphate dehydrogenase (GAPDH) was used as the housekeeping gene. Reverse transcriptase control (–RT) was used in all the experiments to monitor genomic DNA contamination.

### 9. Estimation of total intracellular reactive oxygen species (ROS) production

The 2′,7′-dichlorofluorescein diacetate (DCFDA; Sigma D6883) was used to estimate intracellular ROS production. Briefly, cells were stained with 10 μg/mL DCFDA dye was incubated for 20 min, followed by washing PBS. Imaging was done using Olympus IX83 fluorescent microscope aided with Cell Sens imaging software at 10x objective magnification. The fluorescence intensity was quantified using ImageJ software. Relative change in fluorescence was expressed in fold change increases compared to the control cells(*24*).

### 10. Trans-Endothelial Electrical Resistance (TEER)

To evaluate the integrity of the *in vitro* BBB model upon CM treatment, we measured permeability values every 12 h up to 5 days. An EVOM2 volt-ohm meter (Merck-Milipore) was used with STX2 electrode sets. Briefly, The TEER of blank media was subtracted from the TEER of all groups and was calculated as,

TEER (Ω/cm^2^) = TEER (Ω) x surface area (0.33 cm^2^)

### 11. Western blot

Directly infected AGS CM-exposed HBEC-5i cells and IMR-32/U-87MG cells were harvested and washed with PBS. The cells were lysed with radioimmunoprecipitation assay buffer (RIPA lysis buffer, VWR, Radnor, USA) as described previously(*25*). Antibodies against ZO-1 (#40-2300, Invitrogen), E-cadherin (ABM43F8, Abgenex), VE-cadherin (14-1449-82, Invitrogen), β-catenin (14903, CST), Vinculin (4650, CST), ATG7 (A0691, Abclonal), GFAP (12389, CST), MBP (13344, CST), and Survivin (2808, CST) were used. GAPDH (2118, CST, USA) was used as a housekeeping gene to test the relative expression. The Gel Documentation system (BioRad, USA) was used to visualize the blots, which were further quantified by ImageJ software (National Institutes of Health, USA).

### 12. Immunocytochemistry (ICC) and Immunohistochemistry (IHC)

Upon completion of the timepoint, cells were fixed with 4% paraformaldehyde for 30 min. Permeabilization was done using 0.2% Triton X-100 for 30 min. Further, blocking was performed using 1% BSA (Sigma, USA). Primary antibodies, namely ZO-1, E-cadherin, VE-cadherin, and Vinculin (1:200). The cells were then washed with 1x PBS and incubated with a mixture of secondary antibody (1:1000) and 4′, 6′-diamidino-2-phenylindole (DAPI). The coverslips were washed and mounted on a slide with an antifade mounting medium. Cells were observed using a fluorescence microscope (FluoView 1000, Olympus, Tokyo, Japan) and quantified using ImageJ software(*23*).

Immunohistochemistry (IHC) protocol was adapted from Abcam and was performed using 5-micron brain tissues sectioned using a microtome. As previously mentioned, the sections were deparaffinized and hydrated in Coplin jars with xylene (5 min), xylene:100% ethanol (3 min), 100% ethanol (5 min), 95% ethanol (3 min), and 70% ethanol (3 min). Finally, sections were washed using 1x PBS and shifted to 10 mM sodium citrate buffer incubated at 80 °C for 2 h in the water bath. After bringing the Coplin jar to room temperature, the sections were washed with 1x PBS and fixed with 4% paraformaldehyde for 30 min, followed by 1x PBS wash. Subsequent steps were similar to the immunocytochemistry protocol. The primary antibodies used were ZO-1, VE-cadherin, GFAP, and MBP.

### 13. Enzyme-Linked Immunosorbent Assay (ELISA)

Sandwich ELISA was performed using a manual from FineTest for TNF-α (EM0183, Fine Test) and NEFL (EM1688, Fine Test) on serum samples of mice. Post-decapitation, mouse blood was collected to separate serum by centrifuging it at 1000g for 20 min at 4 °C. As per the manual, all standard proteins were prepared, and 100 µL of serum was loaded to each well. Tissue samples were diluted with sample buffer to a final concentration of 0.3 mg/mL. Further, the plates were incubated for 90 min at 37 °C. Subsequently, the plates were washed with a washing buffer. A 100 µL of biotin-labeled antibody was added to each well with 1:100 dilution and incubated at 37 °C. After removing the antibody, the plate was washed thrice, and a 1:100 SABC solution was added to each well for 30 min at 37 °C. After washing, 90 µL of TMB solution was added to each well for 20 min. Finally, a stop solution (50 µL) was added, and absorbance was recorded at 450 nm using a microplate reader. The data was normalized using control, and protein concentration in pg/mL or ng/mL was calculated using standard curve.

### 14. Haematoxylin and Eosin staining

After the treatments, a portion of the mice brain was fixed with formalin and stored at 4°C. Paraffin-embedded sections were subjected to microtome to achieve sections of 4µm thickness and were further utilized for haematoxylin and eosin (H&E) staining. Post-paraffinization, sections were subjected to deparaffinization in a coupling jar with xylene for 5 min, followed by incubation in 100% ethanol for 3 min. Subsequent incubations were performed in 95%, 70%, and 50% ethanol, after which the slides were washed with 1x PBS. Nuclear staining was performed by incubating slides in haematoxylin for 5 min, and excess dye was removed with 1% acid alcohol solution (1% HCl in 70% ethanol) for 30 s and washed in tap water. The sections were counterstained with 1% eosin for 1 min and washed in tap water. The slides were air-dried, and sections were mounted using mounting media and observed under a bright-field microscope. The H&E sections of brain and gut tissues were analyzed by a pathologist to classify alterations upon CM treatment.

### 15. Congo red staining

The staining was performed on sagittal brain and gut sections with 4µm thickness. Briefly, the sections were deparaffinized using 3 changes of xylene (2 min) and subjected to sequential rehydration in absolute, 90%, and 70% alcohol in Coplin jars for 2 min each. The hydrated slides were then transferred to 50% alcohol for 30 s and then 50% alcohol + 0.01% sodium hydroxide for 5 min. This was followed by 0.5% Congo red solution and then subjected to a series of dehydration for 2 min each. Finally, the slides were rinsed in xylene for 3 min and mounted with DPX medium(*26*).

### 16. Statistical analysis

All *in vitro* experiments were performed in three biological triplicates. Data were presented as ± standard error mean (SEM) of three data points. Statistical analyses were performed, and graphs were prepared using GraphPad Prism 8 software. Statistical significance was assessed using the student’s unpaired t-test and two-way ANOVA followed by post-hoc analysis. The threshold for significance was established at 5%, aligning with a 95% confidence interval. When p-values were under 0.05, 0.01, and 0.001, they were marked as statistically significant. Increased expression levels were indicated with *, **, and *** symbols, while reduced expression levels were denoted by #, ##, and ### symbols, respectively.

## Results

### 1. Coinfection of *Helicobacter pylori* and Epstein-Barr virus synergistically disrupts gut barrier integrity and cell polarity in gastric epithelial cells

Gastric epithelial cells (AGS) were subjected to *H. pylori*, EBV, or coinfection of both pathogens to observe distinct alterations in gut barrier function (**Figure 1A**). Of note, while both pathogens efficiently replicate in AGS cells, the co-infection did not significantly influence their respective replication levels (**Suppl Figure S1**). The transcript analysis revealed significant upregulation of inflammatory genes such as *IL-8* (p<0.05) along with downregulation of junctional markers, *ZO-1,* and *CDH1* (p<0.05 and 0.001) in the coinfected samples (**Figure 1B**). Cell polarity markers, *Par1,* and *Crumbs* were markedly downregulated (p<0.001) in the coinfected gastric cells (**Figure 1B**). Western blot analysis also revealed a reduced protein expression of junctional and cytoskeletal markers, ZO-1, E-cadherin, Vinculin, and β-catenin (p<0.001, 0.01, 0.05) in coinfected gastric cells (**Figure 1C**). To evaluate the potential impact of these TJ/AJ proteins down-regulation, we measured the trans-epithelial resistance of a tight monolayer of AGS cells seeded in transwell. The results showed that there was a significant disruption in gastric cell junctions in the co-infected conditions (p<0.05) at 30 h (**Figure 1D**), indicative of increased permeability of the gastric cells.

**Figure 1.**
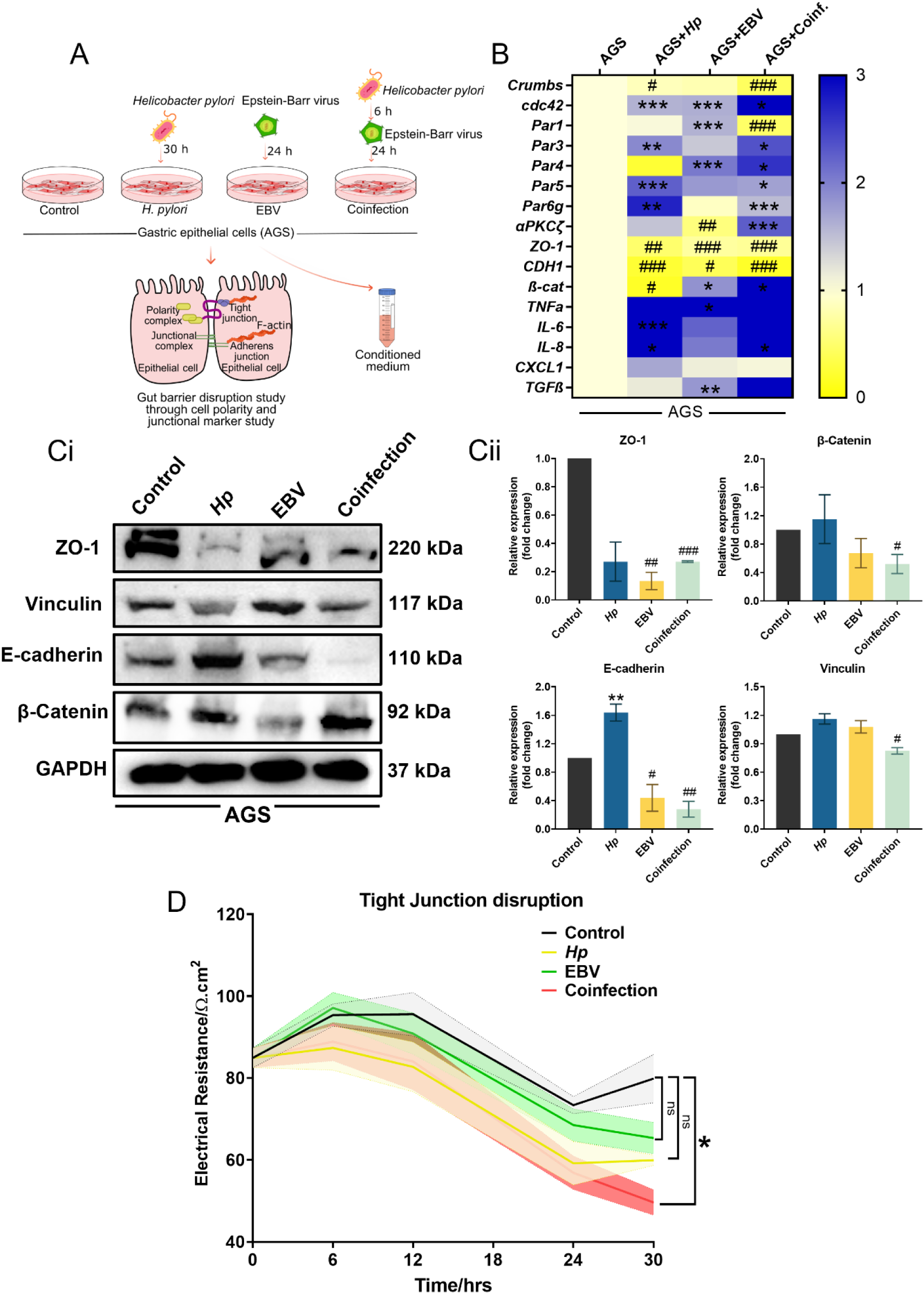
Status of gastric epithelial cell modulation upon infection of *H. pylori*, EBV, and coinfection. (Ai) Infection methodology adapted for infection and coinfection to AGS cells. (Bi) Transcript level alteration of host polarity and junctional markers. (Bii) Infection transcripts modulation in EBV and *H. pylori* genes. (Ci) Representative western blot images indicating a junctional disruption in infected and coinfected cells. (Cii) Quantitative analysis of western blots. (D) Tight junction disruption in gastric epithelia upon coinfection. The p-values of <0.05, <0.01, and <0.0001 are considered statistically significant (unpaired t-tests for qRT-PCR, western blotting, and immunostaining/ two-way ANOVA for TEER) and represented with */#, **/##, and ***/### upregulation/downregulation, respectively.

We also observed similar modulations in immunofluorescence of ZO-1, E-cadherin, and Vinculin (p<0.01 and 0.05) (**Figure 2A and 2B**). Additionally, actin fibres were stained with phalloidin to assess cytoskeletal involvement upon coinfection to gastric cells. In coinfected samples, the cells exhibited an altered morphology (intensity change, p<0.001), indicating the invasive nature of both pathogens to alter gastric epithelium (**Figure 2C**).

**Figure 2.**
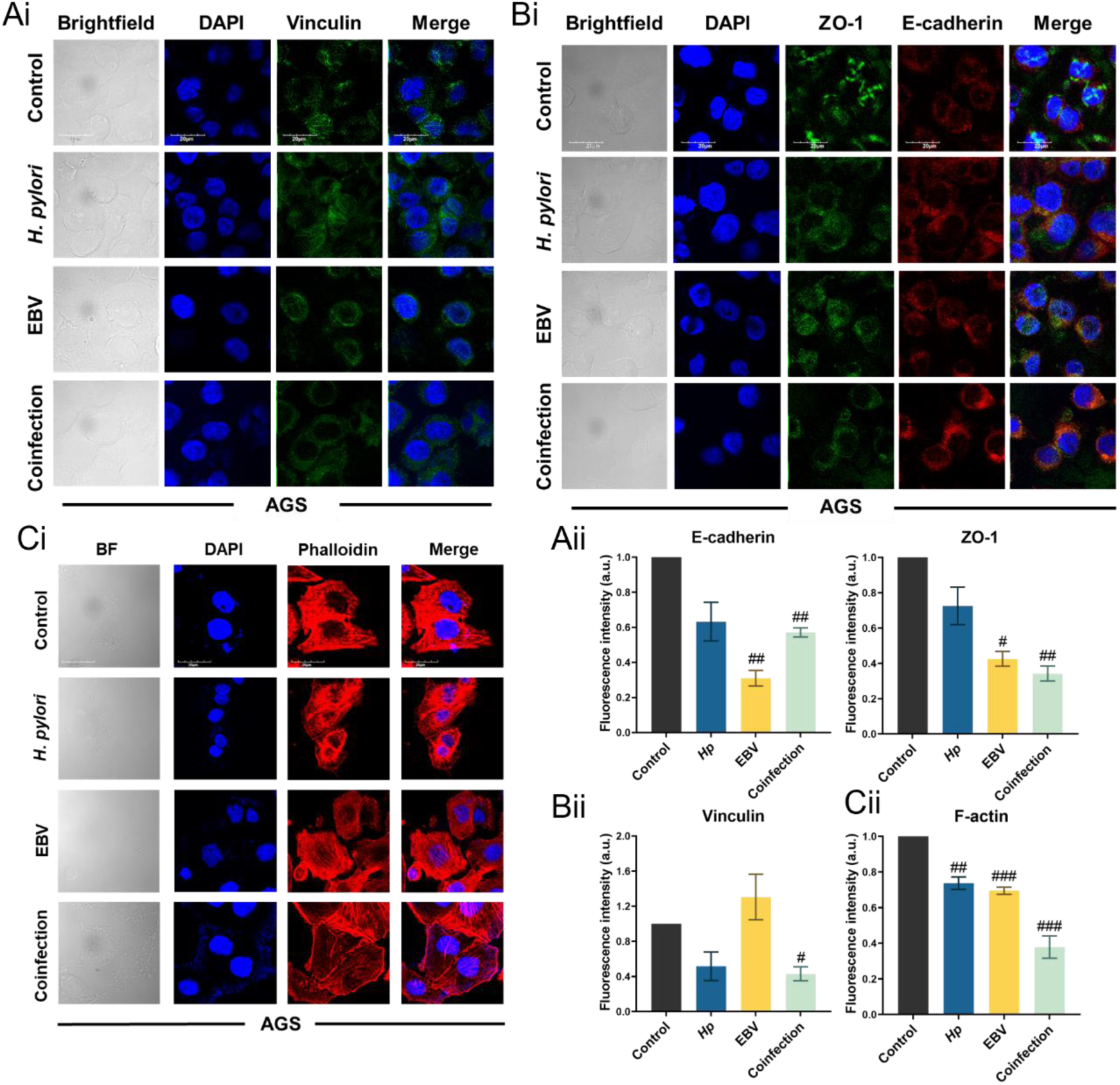
Alteration in junctional markers in infected gastric cells. (Ai) Immunostaining of ZO-1 and E-cadherin revealed the downregulation of both junctional markers in *H. pylori*, EBV, and coinfected cells. (Aii) Quantitative analysis of immunostaining. (Bi) Immunofluorescence of Vinculin revealed a disruptive cytoskeleton. (Bii) Quantification of Vinculin. (Ci) The F-actin cytoskeleton was disrupted severely in the coinfected cells. (Cii) Quantification of F-actin intensity. The p-values of <0.05, <0.01, and <0.0001 are considered statistically significant (unpaired t-tests) and represented with */#, **/##, and ***/### upregulation/downregulation, respectively.

### 2. Acute and chronic systemic inflammation due to gut microbiome disruption aligns with BBB disruption

Dysbiotic gut secretions has been previously shown to have important consequences onto brain functions. Peripheral infections can lead to increased permeability of the BBB. We first analyzed the TJ and AJ proteins to characterize the brain endothelial barrier. As a component of TJ proteins, we analyzed cytosolic ZO-1 together with components of AJs such as VE-cadherin and junctional β-catenin. The BECs were exposed to conditioned media (CM, referring to the secretome of AGS cells) for one and five days (**Figure 3A**). Our results revealed a significant increase in ROS production upon exposure of BEC to CM from co-infected cells, as indicated by intense DCFDA staining (**Figure 3B-C**). Analysis of mRNA expression levels by RT-qPCR revealed significant modulation of AJ/TJ genes in the various CM conditions (**Figure 3D**). First, compared to AGS CM-exposed cells, *Claudin 5* and *Cdc42* were significantly downregulated (p<0.05) in *Hp*CM-exposed brain endothelial cells at D1. Cell polarity markers such as *Par 3, –4, –5,* and *-6* were also downregulated (p<0.01 and 0.001) in *Hp*CM-exposed cells. On the other hand, we observed significant upregulation (p<0.05 and 0.01) of *VEGFA, Claudin 5, Cx43*, and *ZO-1,* while downregulation (p<0.05 and 0.01) of *CDH5, Par1, –4, and –5* indicating abrupt expression of polarity and junctional markers upon exposure to *H. pylori*-infected gastric cell conditioned medium indicating dyshomeostasis and mislocalization. At D1, EBV CM exposed cells showed significant downregulation (p<0.05 and 0.01) of *Integrin β3, Occludin, Par4* and *Par6g,* while at D5, *Par1,* and *Par5* were significantly downregulated (p<0.05). Interestingly, upon exposure to coinfected CM, significant downregulation (p<0.05 and 0.01) was observed in *VEGFA, Claudin5, Cx37, cdc42, Par1*, and *Par6g* while a significant upregulation (p<0.05 and 0.01) was observed in *ZO-1, CDH5*, and *Par4*. Meanwhile, at D5, we observed an upregulation (p<0.05 and 0.01) of *Integrin β3, β5, VCAM1, Claudin5, Cx43, Cx37,* and *Par4* while a significant downregulation (p<0.05 and 0.001) of *Integrin β1, VEGFA, Cx40,* and *CDH5* was revealed (**Figure 3D**). At the protein level, we observed significant upregulation of ZO-1 (p<0.01) while a significant downregulation in β-catenin (p<0.0.5) at D1 of coinfected CM-exposed HBEC-5i cells. In contrast to D1, at D5, the levels ZO-1 (p<ns), VE-cadherin (p<ns), and β-catenin (p<0.001) were downregulated (**Figure 3E-F**). A similar expression pattern was observed through immunostained HBEC-5i cells with significant downregulation of ZO-1 at both D1 and D5 (p<0.05 and 0.01) (**Suppl Figure S2**).

**Figure 3.**
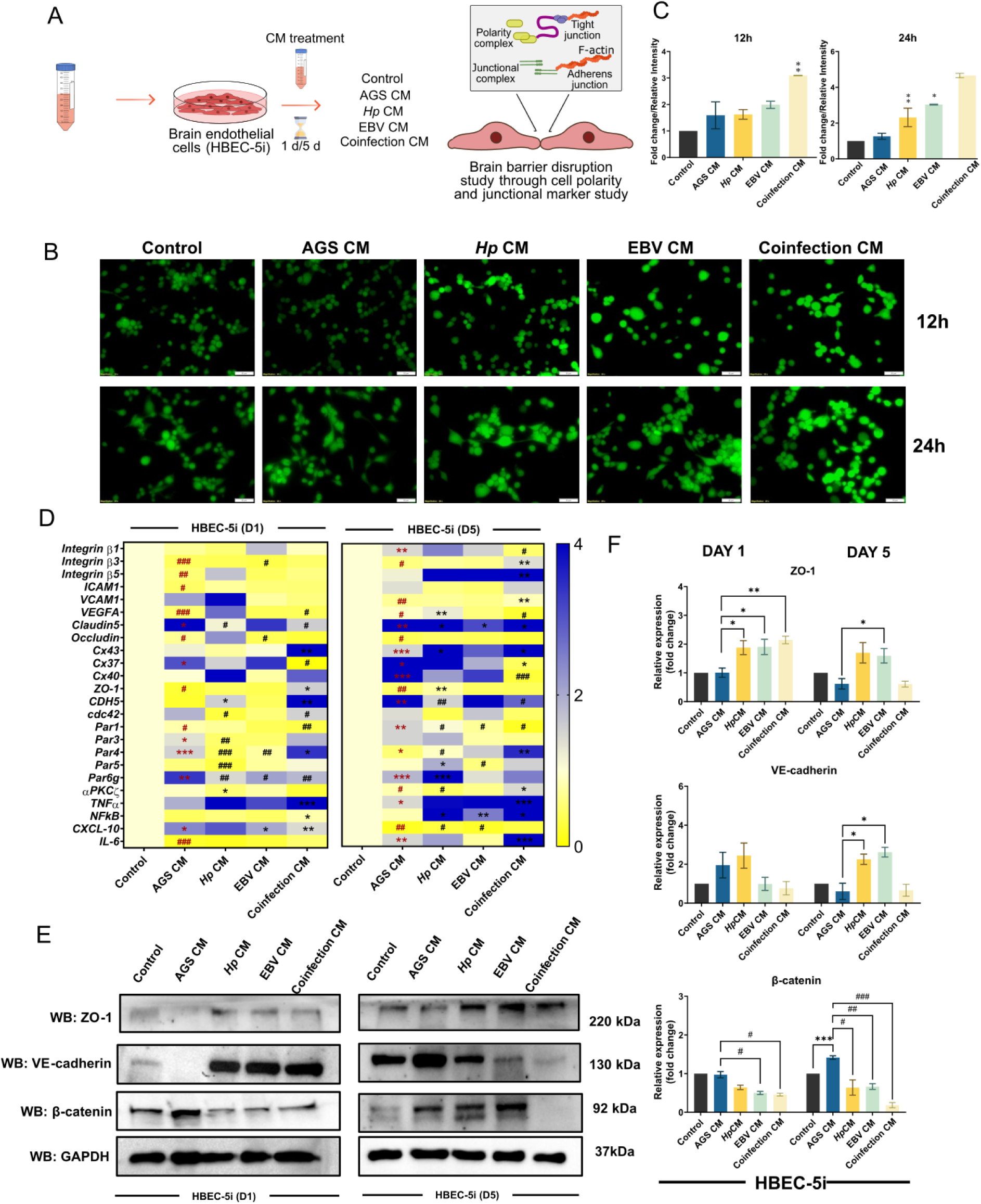
Disruption of brain endothelia upon exposure to infected secretome. (A) Methodology of conditioned media exposure. (B) ROS induction at early time points upon secretome exposure. (C) Quantification of DCFDA staining. (D) Transcript modulation of polarity and junctional markers at D1 and D5 of secretome exposure to brain endothelial cells. (E) Western blotting shows altered expression of junctional markers at D1 and D5. (F) Quantification of western blots. The p-values of <0.05, <0.01, and <0.0001 are considered statistically significant (unpaired t-tests) and represented with */#, **/##, and ***/### upregulation/downregulation, respectively.

### 3. Infected secretome causes blood-brain barrier dysfunction associated with neurological damage

The loss of integrity of the NVU represents a major event associated to neurodegeneration. We adapted a transwell methodology to assess how interference with BBB endothelial cells affect the neurological milieu. Specifically, we evaluated the impact of CM-exposed endothelial cells on co-culture of neurons and astrocytes (**Figure 4A**). Messenger RNA expression levels of the neural co-culture were measured by RT-qPCR and revealed increased *GFAP* and *BDNF* gene expression at D1 (p<0.05) upon exposure to all CM conditions (**Figure 4B**). In contrast, a downregulation (p<0.05) of *NLRP3* was observed in *Hp-*CM-exposed cells, while an upregulated and downregulated expression pattern was observed in *GFAP* and *LC3B*, *BDNF,* and *PSPN,* respectively. EBV-CM exposure led to an increase in expression (p<0.05 and 0.01) of *Survivin, GFAP*, and *PSPN*, while a decreased expression (p<0.05) of *MBP* at D1. At D5 in this condition however, we observed an upregulation (p<0.05) of *GFAP, MBP, LRP1*, and *PSPN*. We noted an increase (p<0.05 and 0.01) in *NLRP3, LC3B, GFAP*, and *PSPN* while declined expression (p<0.05 and 0.01) of *MBP, LRP1*, and *BDNF* upon exposure of coinfected-CM for D1. Of note, when the coinfected CM was exposed for 5 days, we observed a heightened expression (p<0.05 and 0.01) of *Survivin* and *GFAP* while a reduced expression (p<0.05, 0.01, and 0.001) of *LC3B, MBP, LRP1, BDNF*, and *PSPN*. Additionally, we also observed significant perturbations in the inflammasome of the co-cultured cells exposed to coinfected CM for up to 5 days with an upregulation (p<0.05 and 0.01) of *NFκB*, *TNFα, IL-6, IL-10, IL-1β,* and *CXCL10* (**Figure 4B**).

**Figure 4.**
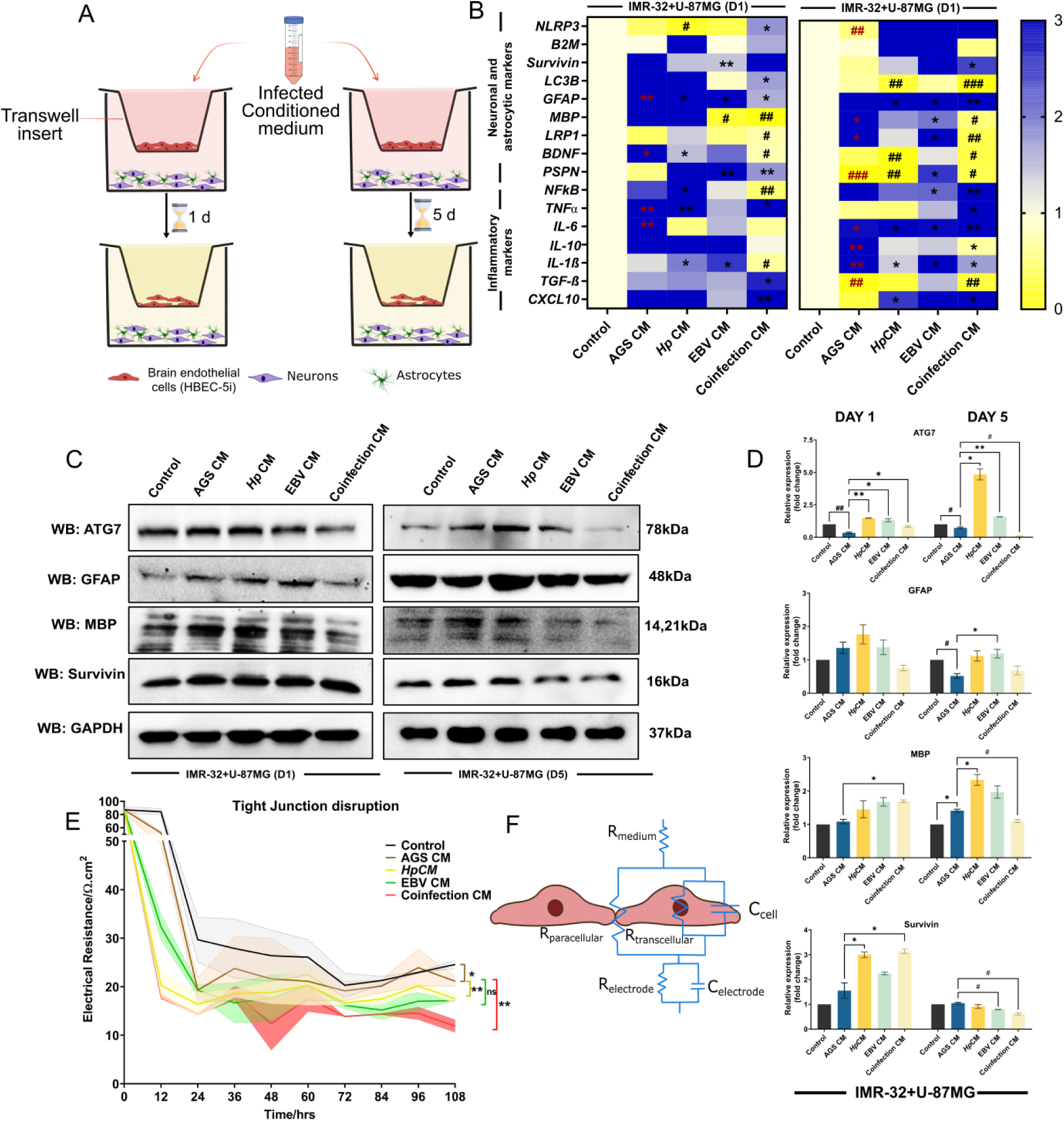
Neurological deficits associated with infected secretome exposure to neuron and astrocyte co-cultures. (A) Transwell methodology to observe BBB disruption and subsequent exposure to the neural milieu. (B) Transcript level modulation of neuronal and astrocytic activation markers at D1 and D5 of CM exposure. (C) Western blotting of junctional proteins in endothelial cells. (D) Quantification of western blot. (E) TEER assay assesses barrier disintegration upon coinfected CM-exposed cells until D5. (F) Demonstration of cellular permeability disintegration. The p-values of <0.05, <0.01, and <0.0001 are considered statistically significant (unpaired t-tests) and represented with */#, **/##, and ***/### upregulation/downregulation, respectively.

These results were further confirmed at the protein level, as we observed a significant downregulated expression of ATG7 (p<0.05) and GFAP while upregulation of MBP (p<0.05) and Survivin (p<0.05) post 1-day exposure of coinfected CM (**Figure 4C-D**). Upon exposure of the same up to D5, we observed a significant downregulation of ATG7, MBP, and Survivin (p<0.05), suggesting altered functional autophagy in neurons and activated glia (**Figure 4C-D**). Finally, we found that these changes correlated with the loss of permeability of the endothelial barrier as measured by TEER in *Hp-*CM (p<0.01), EBV-CM (p<ns), and Coinfection-CM (p<0.01) in comparison to control CM (**Figure 4E-F**).

### 5. Sustained inflammation through secretome exposure indicates BBB disruption *in vivo*

Dysbiosis in gut microbiota can induce BBB breakdown *in vivo*(*27*). This was validated by assessing the expression of ZO-1 and VE-cadherin by confocal microscopy. The TJ and AJ morphology were assessed in the hippocampal and cortical regions surrounding the same. A strong and continuous cerebrovasculature was normal in control mice’s cortex and hippocampus (**Figure 5A**). Compared to controls, coinfected CM-exposed mice exhibited punctate staining of both proteins, indicating abnormal morphology of the blood vessels. This was evident through immunohistochemistry of ZO-1 and VE-cadherin (p<0.05) indicating significant downregulation in the coinfected CM-exposed mice (**Figure 5B**).

**Figure 5.**
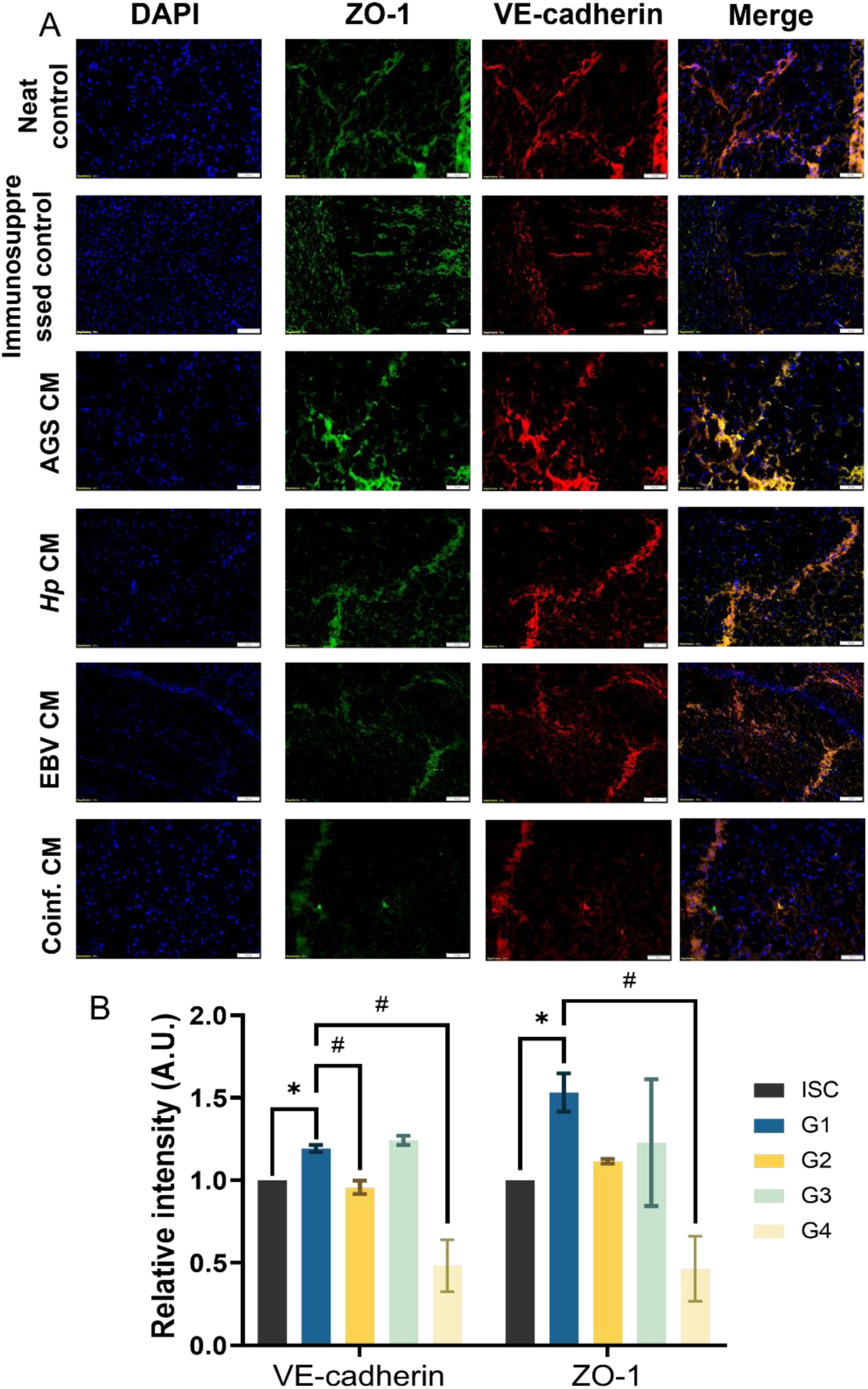
Alteration in junctional markers in mice cortex. (A) Immunostaining of ZO-1 and E-cadherin. (B) Quantitative analysis of immunostaining. The p-values of <0.05, <0.01, and <0.0001 are considered statistically significant (unpaired t-tests) and represented with */#, **/##, and ***/### upregulation/downregulation, respectively. (G1: AGS CM exposed mice, G2: Hp CM exposed mice, G3; EBV CM exposed mice, G4: Coinf. CM exposed mice).

In line with our *in vitro* results, we observed an enhanced expression of GFAP (p<0.05) in coinfected CM mice brain tissue sections, indicating glial activation in the cerebral compartment (**Suppl Figure S3**). Contrarily, immunohistochemistry of MBP revealed its enhanced expression (p<0.05), indicating possible progression towards multiple sclerosis through myelin toxicity *in vivo*.

### 6. CM-exposed mice exhibit stress vulnerability and neurological deficit

The TJs of the endothelium selectively seal the BBB to prevent viral neuroinvasion(*28*). However, pathogens are associated to BBB dysfunctions, allowing viruses to cross the BBB in various ways to reach the CNS(*29*). Perturbation of BBB integrity has been associated to neurodegenerative disorders(*30*), the latter being well correlated to virus exposure(*31*). Here we tested the effect of CM on neurodegeneration, by staining amyloid plaques using Congo red (**Figure 6A-B**). We found increased level of amyloid plaques in the hippocampal region of mice exposed to *Hp-*CM (p<0.01), EBV-CM (p<0.05), and coinfected CM (p<0.01). Moreover, increased inflammatory responses were observed in all mice groups. Specifically, we observed an elevation of TNF-α in the serum upon treatment with coinfected-CM. Neurofilament L (NfL) are cytoskeletal proteins that shed upon injury in the nervous tissue and thus become prominent markers for neurological diseases since neurofilaments can also diffuse across the BBB. We found a significant increase in serum NfL levels of CM-exposed mice (p<0.05) (**Figure 6C-D**).

**Figure 6.**
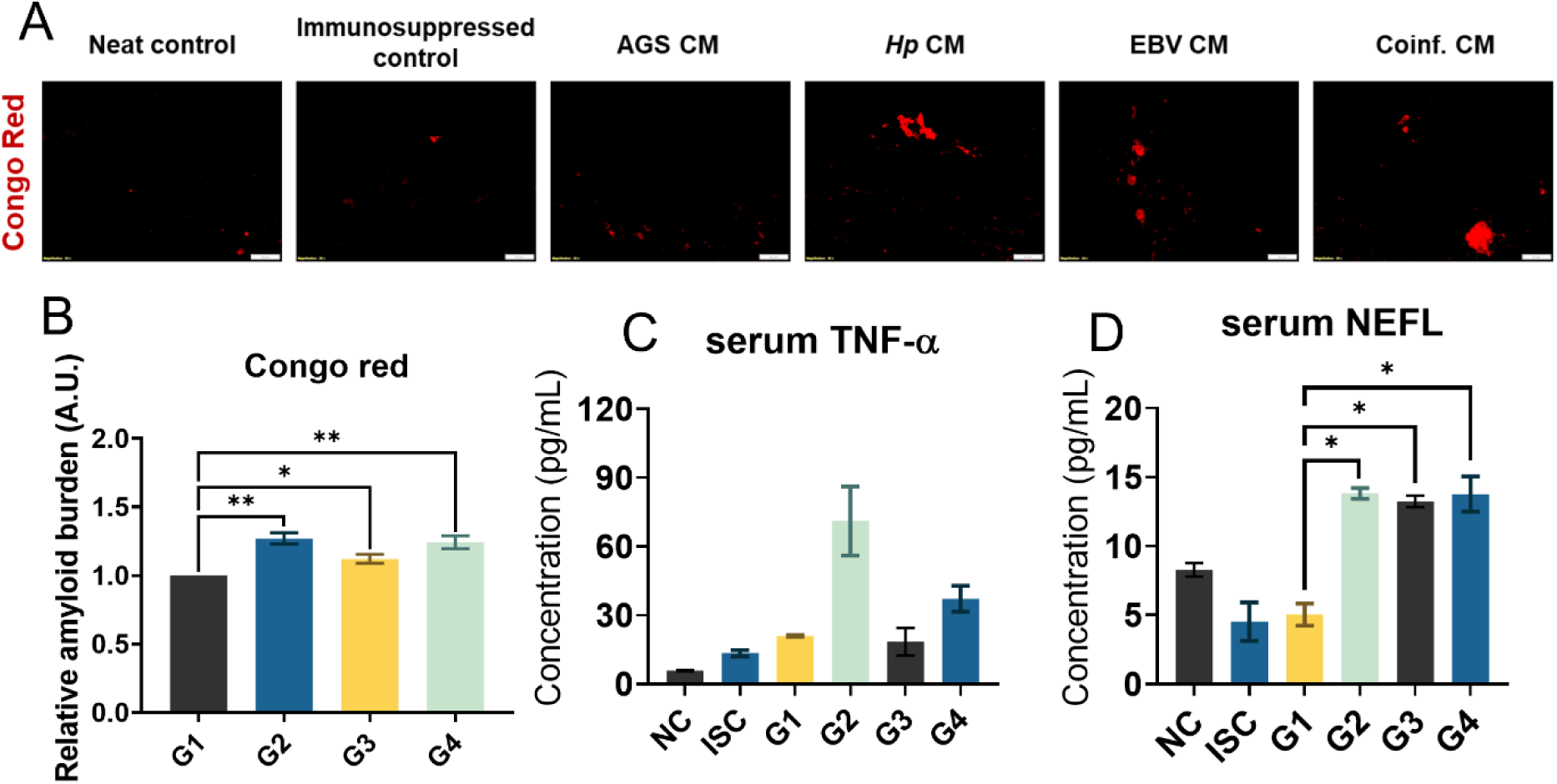
Amyloidogenic plaques and enhanced presence of secretory markers were observed in coinfected CM-exposed mice through Congo Red staining and ELISA, respectively. (A) Congo Red staining of mice cortex. (B) Quantitative analysis of staining. (C, D) Serum concentrations of TNF-α and NEFL in mice serum detected through ELISA. The p-values of <0.05, <0.01, and <0.0001 are considered statistically significant (unpaired t-tests) and represented with */#, **/##, and ***/### upregulation/downregulation, respectively.

In order to assess whether the observed amyloid plaques were associated with marked neurocognitive impairment, we the performed a behavioural experiment using the the 8-arm radial maze assay (**Figure 7A**). The dentate gyrus region of mice hippocampus is composed of granular cells with compact cellular zones consisting of neuronal processes. The noticeable thick bundles of cells in hippocampus were shrunk in infected secretome exposed mice with disrupted arrangement of nuclei (**Figure 7B**). We observed that mice exhibited impaired memory at 5 weeks post exposure to coinfected CM, including a lower reference (p<0.05) and working memory (p<0.05) functionality when compared to AGS CM as mice took longer to find food baits (**Figure 7C**). Finally, upon subjecting mice to an elevated-plus maze, we observed the development of anxiety-like symptoms in mice (**Figure 7D**). This was observed due to the lack of mobility of mice in open arms and their preference for closed arms upon exposure to coinfected CM.

**Figure 7.**
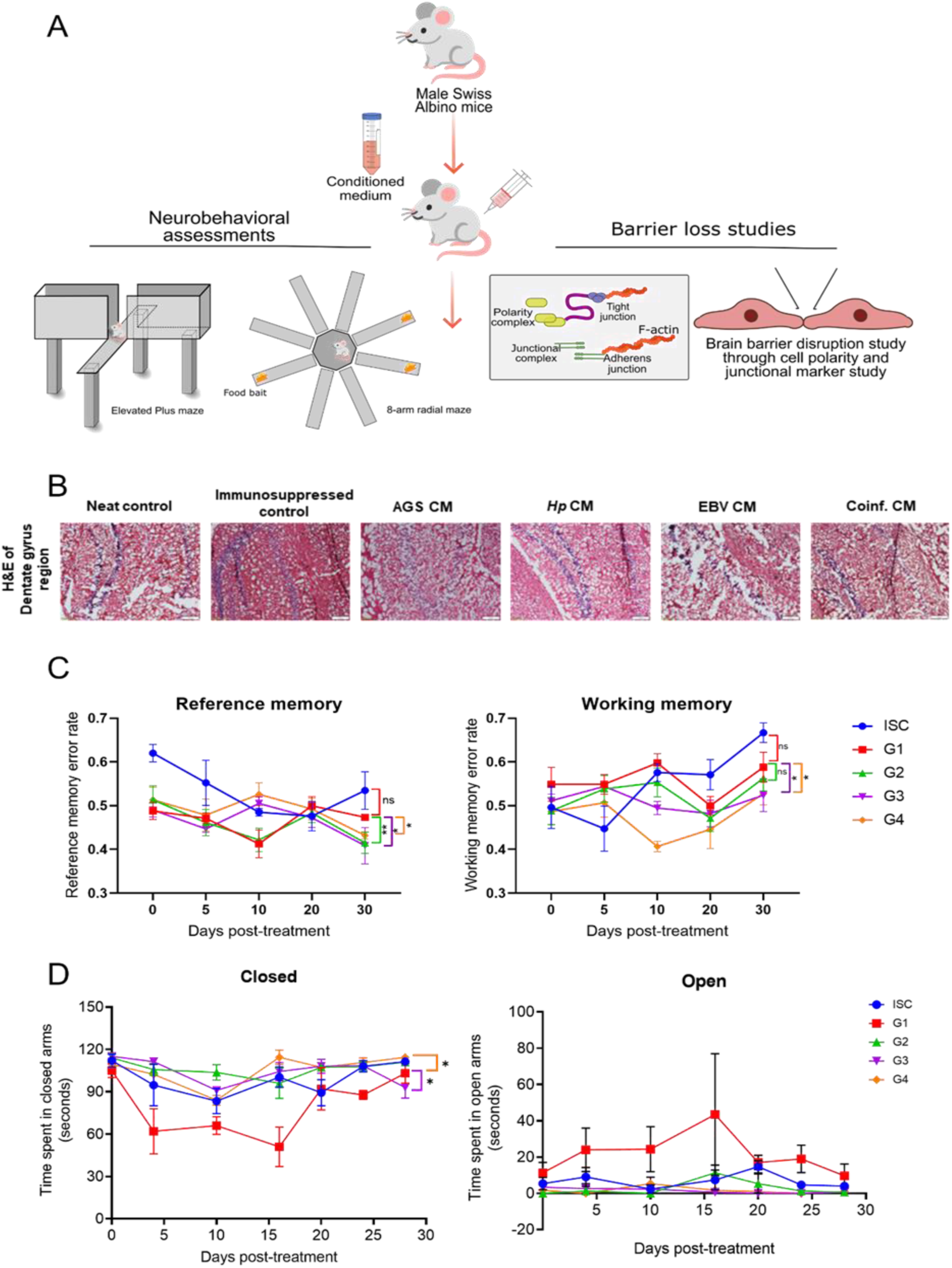
*In vivo* methodology and neurobehavioral alterations in mice. (A) Methodology of secretome exposure to mice. (B) H&E staining of mice groups. (C) Reference and working memory changes were observed during the radial-arm maze test. (D) Analysis of anxiety-like behaviour through elevated-plus maze test. The p-values of <0.05, <0.01, and <0.0001 are considered statistically significant (one-way ANOVA) and represented with */#, **/##, and ***/### upregulation/downregulation, respectively.

**Figure 8.**
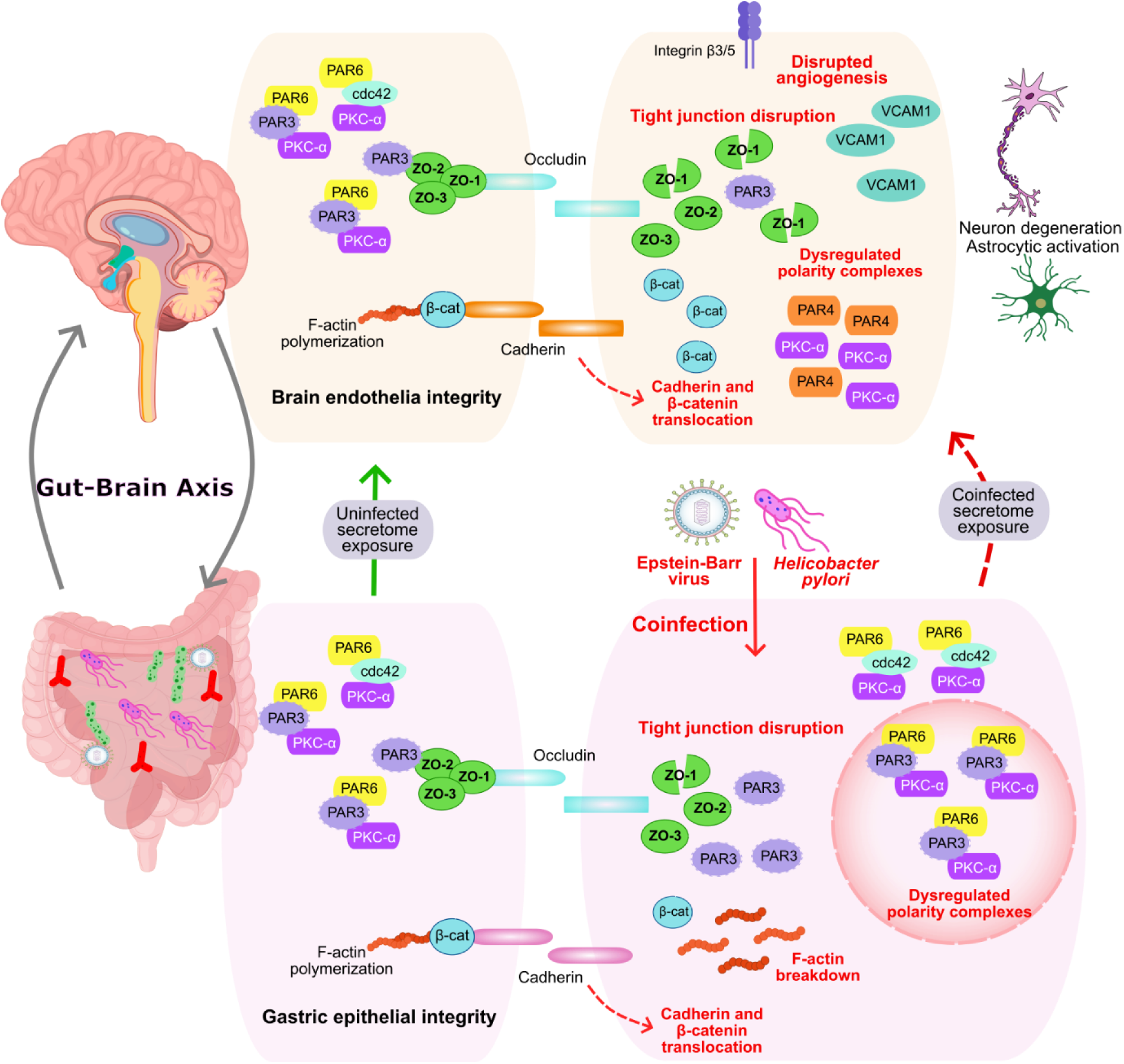
A representation of molecular pathways disrupted in gut-brain axis due to coinfection of *Helicobacter pylori* and Epstein-Barr virus to gastric epithelial cells. Healthy tissues (left panels) exhibit intact barriers maintained by organized protein complexes including polarity regulators and junction components, preserving normal gut-brain communication. Pathogen coinfection (right panels) induces barrier compromise through disorganization of these protein networks, causing F-actin fragmentation and junction protein mislocalization. The resulting barrier dysfunction enables microbial secretome penetration, triggering pathological consequences including aberrant blood vessel formation, elevated VCAM1 expression, and neural tissue damage (astrocyte reactivity and neuronal death). This molecular mechanism illustrates how enteric infections may contribute to neurological disorders through disruption of epithelial and endothelial boundaries.

## Discussion

Reduced gut and brain barrier integrity precedes a disrupted gut-brain axis likely to be a causal factor or a helping agent to exacerbate neurological deterioration. The main findings of this study were to elucidate the mislocalizations of cadherins and occludins in maintaining the gastrointestinal barrier and blood-brain barrier in coinfections mimicking gut dysbiosis. Several investigators have utilized infections of gut pathogens such as *Clostridium perfringens* to the intestinal epithelial monolayers to demonstrate barrier dysfunction(*32*). *H. pylori* is one such member of the gut microbial community, which, once provided conducive environment, leads to an increase in gastric pH and induces its effects through characteristic proteins such as CagA, VacA, BabA, OipA, etc(*7*). Beyond the gut microbiome, the gut virome also plays a crucial role in maintaining homeostasis. Several reports have indicated a high prevalence of EBV as a part of gut virome eliciting severe immunogenic responses in the host(*33*, *34*). Dysbiosis to both microbiome and virome can lead to severe manifestations. Due to the crucial role of gut microbiota in the gut-brain axis, modifications in gastrointestinal motility, heightened permeability, and a weakened defense mechanism can cause ‘second-brain aging(*35*).’ Secretory factors such as short-chain fatty acids (SCFAs) are modulated upon *H. pylori* infection, disrupting gut homeostasis. As metabolites can affect the transport and epigenetic machinery of the cells, they can also lead to EBV reactivation, thus inducing pathogenicity in distant organs.

Here, we examined the hypotheses that gut leakage and subsequent transport of leaked moieties to BBB increased upon coinfection due to the dysfunction of TJ/AJ proteins. Gut hyperpermeability imitating a leaky gut due to coinfection of *H. pylori* and EBV disrupts the TJ/AJ complexes. Interestingly, we also observed that coinfection led to the remodelling of cell polarity markers and actin cytoskeleton in gastric cells. *H. pylori* and EBV are well known to usurp cell polarity(*17*, *36*). Loss of ZO-1 and E-cadherin in our study revealed their vulnerability to coinfections. When connected with disruption of polarity and cytoskeleton, this points out to a severe complication(*37*). One of the most studied sites of gut dysbiosis is the central nervous system (CNS), inducing neurological malformations. A disrupted gut-brain axis leads to neurological manifestations. However unknown, several mechanisms have been pointed out to elucidate the role of gut microbiota in neurological impairment, as also seen in our previous studies indicated by altered JAK-STAT signaling mechanisms(*38*).

The interaction of viruses and microbes is a common phenomenon in the host and is understudied in aspects of the gut-brain axis. We attempted to expose the secretome of coinfected gastric cells and assessed the disruptions in BBB. BBB hyperpermeability occurs due to loss of interaction among brain endothelial cells and is postulated to be dangerous, ultimately promoting AD pathogenesis(*39*). BBB dysregulation was observed up to day 5 of coinfected secretome exposed samples compared to alone infections as evidenced by loss of VE-cadherin, β-catenin, and ZO-1. It is possible that the internalization of VE-cadherin led to increased permeability in BECs. Upregulation of Par4 and VCAM1 is linked to dysregulated angiogenesis and inflammation in endothelial cells which was also seen in our coinfection secretome exposed cells(*40*). Our findings suggest that the leaky barrier also led to neuronal death and glial activation in a BBB mimic model indicated by co-culture of astrocytes and neurons. We observed that autophagic flux was disrupted (ATG7), essential for neuronal homeostasis. Cell survival mechanism was also altered, indicated by Survivin downregulation inducing stress in neurons and glia. Astrogliosis indicated by GFAP upregulation and myelin dysregulation indicated by MBP aligns with potential neurodegenerative changes. Survivin downregulation was correlated with decreased neurogenesis(*41*).

The involvement of junctional dysregulation in the formation of leaky barrier thus becomes a crucial point in studying infectious disease, particularly neurodegeneration through gut-brain axis dysregulation(*42*). Long-term exacerbations of coinfected secretome exposure were studied *in vivo*. Herein, we identified systemic inflammation was induced due to coinfected secretome exposure as shown by IL-6 in ELISA of serum samples. Yet another characteristic marker, NfL, was enhanced in serum indicative of neuroaxonal damage secondary to BBB leakage as shown in several other diseases such as cerebral malaria and multiple sclerosis(*43*). We also observed loss of VE-cadherin and ZO-1 from cerebral microvessels in the cortical brain regions of mice, indicative of BBB leakage. Impairments to BBB might lead to neurological dysfunction. A leaky barrier might be correlated to the activation of astrocytes through GFAP and amyloid deposits in the brain, worsening neurological functions in mice. A negative regulation of MBP was observed in coinfected secretome exposure, suggesting possible demyelination. As leaky barriers expose the otherwise separated cerebral microenvironment, a massive flow of cytokines and pathogenic secretions could induce such losses to gut and brain barriers, ultimately causing disruptions in the homeostatic gut-brain axis.

In conclusion, negative regulations in junctional and polarity proteins could be characteristics of pathogenic mechanisms of *H. pylori* and EBV coinfection. Our findings show that a dysfunctional brain microenvironment would thus become imperative to treat as these pathogens become major contributors to neurological inflammation and dysfunction. The study becomes crucial as the gut-brain axis is the primary coordination centre through the exchange of peptides, hormones, immune secretions, and metabolites. A disrupted epithelial barrier would lead to leaky gut formation, becoming an easy route for traversing of pathogenic metabolites to cross the gut and reach the CNS microenvironment. Further, as we observed a deficit in BBB integrity, it becomes an opportunistic moment for the metabolites to induce neurological dysfunction after crossing the BBB. To what extent the coinfection leads to pathogenesis remains elusive. Moreover, considering our current observations, a deeper understanding of signaling pathways inducing such pathogenesis is yet another aspect of the future studies. The neuroplastic mechanisms of such coinfections intervening pathways of calcium excitotoxicity also become important domains to study(*44*, *45*). Moreover, the specific pathogenic metabolites can be targeted to treat gut dysbiosis and associated neurological manifestations.

## Acknowledgments

We acknowledge the Indian Institute of Technology Indore for providing facilities and support and thank lab colleagues for their meaningful discussions. We appreciate Sophisticated Instrument Centre at IIT Indore for providing confocal microscopy facility. We also thank the Ministry of Education and University Grants Commission for the fellowships of Vaishali Saini (PMRF ID: 2103365) and Siddharth Singh, Buddhadeb Baral, and Pramesh Sinha, respectively. We also acknowledge DST-FIST support project No. SR/FST/LS-I/2020/621 for providing us instrumentation facility. We are thankful to Dr. Lilly Ganju and Dr. Neha Jaiswal (Indore and Index Medical College, Indore) for providing the facility of microtomy.

## Contributions

*Conceptualization*: Hem Chandra Jha and Vaishali Saini; *Methodology*: Vaishali Saini and Budhadev Baral; *Experimentation*: Vaishali Saini, Siddharth Singh, Anamika Singh, Pramesh Sinha; *Writing – original draft preparation*: Vaishali Saini; *Writing – review and editing*: Hamendra Singh Parmar, Raphael Gaudin and Hem Chandra Jha; *Writing – preparation of figures*: Vaishali Saini, Siddharth Singh; *Funding acquisition*: Hem Chandra Jha; *Supervision*: Hem Chandra Jha.

